# Economic evaluation of a healthy lifestyle intervention for chronic low back pain: a randomised controlled trial

**DOI:** 10.1101/296285

**Authors:** A Williams, JM van Dongen, SJ Kamper, KM O’Brien, L Wolfenden, SL Yoong, RK Hodder, H Lee, EK Robson, R Haskins, C Rissel, J Wiggers, CM Williams

**Affiliations:** School of Medicine and Public Health, Hunter Medical Research Institute, University of Newcastle, Newcastle NSW, 2308, Australia; Hunter New England Population Health, Locked Bag 10, Wallsend NSW, 2287, Australia.; Centre for Pain, Health and Lifestyle, NSW, Australia; Department of Health Sciences, Faculty of Science, Vrije Universiteit Amsterdam, Amsterdam Public Health research institute, the Netherlands; Department of Health Sciences, Faculty of Science, Vrije Universiteit Amsterdam, MOVE research institute Amsterdam, the Netherlands; School of Public Health, University of Sydney, Lvl 10, King George V Building, Camperdown NSW, 2050, Australia; Neuroscience Research Australia (NeuRA), PO Box 1170, Randwick NSW, 2031, Australia; Centre for Statistics in Medicine, Nuffield Department of Orthopaedics Rheumatology and Musculoskeletal Sciences, University of Oxford, Oxford, UK; Outpatient Services, John Hunter Hospital, Hunter New England Local Health District, Locked Bag 1, New Lambton NSW, 2305, Australia; NSW Office of Preventive Health, Liverpool Hospital, South West Sydney Local Health District, Locked Bag 7279, Liverpool BC 1871

**Author notes:** **Corresponding author:** Amanda Williams.

## Abstract

We performed an economic evaluation of a healthy lifestyle intervention targeting weight loss, physical activity and diet for patients with chronic low back pain, who are overweight or obese. Eligible patients with chronic low back pain (n=160) were randomised to an intervention or usual care control group. The intervention included brief advice, a clinical consultation and referral to a 6-month telephone-based healthy lifestyle coaching service. The primary outcome was quality-adjusted life years (QALYs). Secondary outcomes were pain intensity, disability, weight, and body mass index. Costs included intervention costs, healthcare utilisation costs and work absenteeism costs. An economic analysis was performed from the societal perspective. Mean total costs were lower in the intervention group than the control group (-$614; 95%CI: -3133 to 255). The intervention group had significantly lower healthcare costs (-$292; 95%CI: -872 to -33), medication costs (-$30; 95%CI: -65 to -4) and absenteeism costs (-$1000; 95%CI: -3573 to -210). For all outcomes, the intervention was on average less expensive and more effective than usual care, and the probability of the intervention being cost-effective compared to usual care was relatively high (i.e. 0.81) at a willingness-to-pay of $0/unit of effect. However, the probability of cost-effectiveness was not as favourable among sensitivity analyses. The healthy lifestyle intervention seems to be cost-effective from the societal perspective. However, variability in the sensitivity analyses indicates caution is needed when interpreting these findings.

## Background

Low back pain places a substantial burden on society. Globally, low back pain is ranked first in terms of disability burden, and sixth in overall disease burden.^1^ Low back pain is also very costly, total annual costs are estimated at $9.2 billion in Australia,^2^ and £11 billion in the United Kingdom,^3^ with the largest proportion of these costs attributed to healthcare service use and lost work productivity.^4^ Given the economic burden of low back pain, undertaking economic evaluations of low back pain management approaches is important.

Systematic reviews show that the development and persistence of low back pain is linked to ‘lifestyle risk factors’, such as overweight and obesity.^5^ Interventions targeting lifestyle changes including weight loss, increasing physical activity and improving diet, present a novel and promising strategy to improve outcomes (e.g. pain or disability) for patients with low back pain. In response to a lack of research in this area,^6,7^ we conducted the first randomised controlled trial (RCT) of a healthy lifestyle intervention for patients with chronic low back pain who are overweight or obese.^8^ The intervention involved brief telephone advice, a clinical consultation and referral to a 6-month telephone-based healthy lifestyle coaching service. The primary goal of the intervention was to reduce pain intensity, by reducing weight and improving physical activity and diet behaviours. The purpose of the current study is to undertake an economic evaluation of the healthy lifestyle intervention, compared with usual care.

Economic analyses can be performed from various perspectives including the societal, and healthcare perspectives.^9^ The societal perspective includes all costs regardless of who pays. This frequently incorporates direct costs; intervention costs, plus costs of care unrelated to the intervention (i.e. healthcare services and medication costs), and the indirect costs; absence from work and impact on productivity.^9,10^ In contrast, the healthcare perspective only includes direct costs i.e. intervention costs and the costs of other care.^9^ In this study the primary analysis was conducted from a societal perspective and a secondary analysis was conducted from the healthcare perspective.

## Methods

### Design

We performed an economic evaluation alongside a two-arm pragmatic parallel group RCT, which was part of a cohort multiple RCT.^11^ The study design is described in detail elsewhere.^8,12^ The trial was prospectively registered with the Australian New Zealand Clinical Trials Registry (ACTRN12615000478516). Ethical approval was obtained from the Hunter New England Human Research Ethics Committee (approval No. 13/12/11/5.18) and the University of Newcastle Human Research Ethics Committee (approval No. H-2015-0043).

### Participants

We invited all patients with chronic low back pain who were on a waiting list for outpatient orthopaedic consultation at the John Hunter Hospital, New South Wales (NSW), Australia, to participate in a cohort study involving telephone assessments. All patients in the cohort were informed that regular surveys were being conducted as part of hospital audit processes and to track patient health while waiting for consultation. During one of the telephone assessments, participants of the cohort study were assessed for eligibility for the RCT. Eligible consenting patients were then randomised to study conditions: i) offered the intervention (intervention group), or ii) remained in the cohort follow-up (usual care control group). Due to the design of the study (i.e. cohort multiple RCT)^11^ participants were not aware of alternate study conditions. Participants from either group remained on the waiting list for orthopaedic specialist consultation and could attend a consultation during the study period if scheduled. Participants were also free to access care outside the study as they saw fit.

Participant inclusion criteria for the RCT were: primary complaint of chronic low back pain defined as: pain between the 12th rib and buttock crease with or without leg pain for longer than 3 months;^13^ average low back pain intensity ≥3 out of 10 on a 0-10 numerical rating scale (NRS) over the past week, or moderate level of interference to activities of daily living (adaptation of item 8 on SF-36); 18 years or older; overweight or obese (body mass index (BMI) ≥27kg/m^2^ and <40kg/m^2^) based on self-reported weight and height; and access to a telephone. Exclusion criteria were: known or suspected serious pathology as the cause of back pain, as diagnosed by their general practitioner (e.g. fracture, cancer, infection, inflammatory arthritis, cauda equina syndrome); previous obesity surgery; currently participating in any prescribed, medically supervised or commercial weight loss program; back surgery in the last 6 months or booked for surgery in the next 6 months; unable to comply with the study protocol that required adaption of meals or exercise due to non-independent living arrangements; any medical or physical impairment precluding safe participation in exercise, such as uncontrolled hypertension; unable to speak and read English sufficiently to complete the study procedures.

### Intervention

Participants randomised to the intervention group were offered an intervention involving brief telephone advice, a clinical consultation with a physiotherapist, and referral to a 6-month telephone-based health coaching service.

Immediately after baseline assessment and randomisation, trained telephone interviewers provided the brief telephone advice. This advice included information that a broad range of factors, including lifestyle risk factors contribute to the experience of low back pain, and description of the potential benefits of weight loss and physical activity for reducing low back pain.

The clinical consultation was a face-to-face consultation (up to one hour) conducted in a community health centre with the study physiotherapist, who was not involved in data collection. As detailed in our protocol,^12^ the consultation was informed by Self Determination Theory and involved two broad approaches; (i) clinical assessment followed by low back pain education and advice, and (ii) behaviour change techniques.^14^

The telephone-based health coaching service was the NSW Get Healthy Information and Coaching Service (GHS).^15^ The service involves 10 individually tailored coaching calls, based on national Healthy Eating and Physical Activity guidelines,^16,17^ delivered over 6 months by qualified health professionals.^15^ The GHS is a telephone-based service to support individuals to modify eating behaviours, increase physical activity, achieve and maintain a healthy weight, and where appropriate includes referral to smoking cessation services.

### Control

Participants randomised to the control group remained on the waiting list for orthopaedic consultation (usual care) and took part in data collection during the study period. No restrictions were placed upon their use of other health services during the study period. Control participants were not aware of the intervention group but were told they would be scheduled a clinical appointment for their back pain in 6 months (i.e. 26 weeks post baseline).

### Measures

The primary outcome for this economic evaluation was quality-adjusted life years (QALYs). Secondary outcomes included pain intensity, disability, weight and BMI. We measured costs in terms of intervention costs, healthcare utilisation costs (healthcare service and medication use) and absenteeism costs due to low back pain. For the primary analysis conducted from the societal perspective, all of these cost categories were included. For the secondary analysis conducted from the healthcare perspective, absenteeism costs were excluded.

### Outcomes

Health-related quality of life was assessed at baseline, 6 and 26 weeks using the 12-item Short Form Health Survey version 2 (SF-12.v2).^18^ The patients’ SF-6D health states were translated into utility scores using the British tariff.^19^ QALYs were calculated by multiplying patients’ utility scores by their time spent in a health state using linear interpolation between measurement points. Back pain intensity was assessed at baseline, 6 and 26 weeks using a 0-10 point NRS. Participants were asked to report the “average pain intensity experienced in their back over the past week”, where 0 was ‘no pain’ and 10 was the ‘worst possible pain’.^20^ Disability was assessed at baseline, 6 and 26 weeks using the Roland Morris Disability Questionnaire (RMDQ).^21^ The RMDQ score ranges from 0 to 24, with higher scores indicating higher disability levels. Self-reported weight (kg) was assessed at baseline, 6 and 26 weeks. BMI was calculated as weight / height squared (kg/m^2^)^22^ using self-reported weight at baseline, 6 and 26 weeks and self-reported height from baseline.

### Cost measures

All costs were converted to Australian dollars 2016 using consumer price indices.^23^ Discounting of costs was not necessary due to the 26-week follow-up.^9^

Intervention costs were micro-costed and included the cost to provide the brief advice, estimated from the development and operational costs of the call and the interviewer wages for the estimated average time (5 minutes) taken to provide the brief advice. Intervention costs also included the cost of a one hour clinical physiotherapy appointment, valued using Australian standard costs.^24^ Lastly, intervention costs included the cost to provide a health coaching call from the GHS multiplied by the number of calls each patient received.^25^ The number of health coaching calls received was reported directly by the GHS.

Healthcare utilisation costs included any healthcare services or medication used for low back pain (other than intervention costs). Healthcare utilisation costs were calculated from a patient reported healthcare utilisation inventory. Participants were asked to recall any health services (the type of services and number of sessions) and medications for their low back pain during the past 6 weeks, at 6 and 26 weeks follow-up. Healthcare services were valued using Australian standard costs and, if unavailable, prices according to professional organisations.^24,26,27^ Medication use was valued using unit prices of the Australian Pharmaceutical Benefits Scheme (PBS)^28^ and, if unavailable, prices were obtained from Australian online pharmacy websites.

The average of the week 6 and week 26 costs per patient was extrapolated, assuming linearity, to estimate the cost over the entire 26-week period.

Absenteeism was assessed by asking patients to report the total number of sickness absence days due to low back pain during the past 6 weeks, at 6 and 26-week follow up. Absenteeism costs were estimated using the Human Capital Approach (HCA),^9^ calculated per patient by multiplying their total number of days off by the national average hourly income for their gender and age according to the Australian Bureau of Statistics.^23^ Absenteeism costs were extrapolated using the same method as described above for healthcare utilisation.

### Statistical analysis

All outcomes and cost measures were analysed under the intention-to-treat principle (i.e. analyses were based on initial group assignment and missing data were imputed). Means and proportions of baseline characteristics were compared between the intervention and control group participants to assess comparability of the groups. Missing data for all outcomes and cost measures were imputed using multiple imputation by chained equations (MICE), stratified by treatment group.^29^ Data were assumed missing at random (MAR). Ten complete datasets needed to be created in order for the loss-of-efficiency to be below the recommended 5%.^29^ We analysed each of the 10 imputed datasets separately as specified below. Following this, pooled estimates from all imputed datasets were calculated using Rubin’s rules, incorporating both within-imputation variability (i.e., uncertainty about the results from one imputed data set) and between-imputation variability (i.e. uncertainty due to missing information).^29^

We calculated unadjusted mean costs and cost differences between groups for total and disaggregated costs (intervention costs, healthcare utilisation costs (healthcare services, medications used) and absenteeism costs). Seemingly unrelated regression (SUR) analyses were performed to estimate total cost differences (∆C) and effect differences for all outcomes (∆E), adjusted for the baseline value of the relevant outcome and potential prognostic factors (baseline pain intensity, time since onset of pain, waiting time for orthopaedic consultation and baseline BMI). An advantage of SUR is that two regression equations (one for ∆C and one ∆E) are modelled simultaneously so that the possible correlation between cost and outcome differences can be accounted for.^30^

We calculated incremental cost-effectiveness ratios (ICERs) for all outcomes by dividing the difference in total costs by the difference in outcomes (∆C/∆E). Uncertainty surrounding the ICERs and 95% confidence intervals (95%CIs) around cost differences were estimated using bias corrected and accelerated bootstrapping (5000 replications). Uncertainty of the ICERs were graphically illustrated by plotting bootstrapped incremental cost-effect pairs on cost-effectiveness planes.^9^ We produced a summary measure of the joint uncertainty of costs and outcomes (i.e. cost-effectiveness acceptability curves [CEACs]) for all outcomes. CEACs express the probability of the intervention being cost-effective in comparison with usual care at different values of willingness-to-pay (i.e. the maximum amount of money decision-makers are willing to pay per unit of effect).^9^ Data were analysed in STATA (v13, Stata Corp).

### Sensitivity analyses

We tested the robustness of the primary analysis, through two sensitivity analyses. First, an analysis was performed excluding one patient with very high absenteeism costs (absenteeism costs > $15,000) (SA1). A second sensitivity analysis involved exclusion of intervention participants who did not have reasonable adherence, defined as not attending the clinical consultation and receiving less than 6 GHS health coaching calls (SA2).

### Secondary analysis

A secondary analysis was performed from the healthcare perspective (i.e. excluding absenteeism costs).

## Results

### Participants

One hundred and sixty patients were randomised into the study (Figure 1). Participant characteristics at baseline are reported in Table 1. At 26 weeks, complete outcome data were available for between 65%-75% of participants, depending on the outcome measure, and 59% of participants had complete cost data at 26 weeks. Thus, 26%-35% of effect measure data and 41% of cost data were imputed (Figure 1).

**Table 1.** Baseline characteristics

| Baseline characteristics | Intervention<br>(n=79) | Control<br>(n=80) |
| --- | --- | --- |
| Gender (male) (n,%) | 31 (39.2) | 34 (42.5) |
| Indigenous status (Aboriginal and/or Torres Strait Islander) (n,%) | 7 (8.9) | 5 (6.3) |
| Country of origin (Australia) (n,%) | 69 (87.3) | 68 (85.0) |
| Employment (n,%) |  |  |
| Employed | 17 (21.5) | 17 (21.3) |
| Unemployed | 15 (19.0) | 9 (11.3) |
| Retired | 27 (34.2) | 29 (36.3) |
| Can't work (health reasons) | 20 (25.3) | 25 (31.3) |
| Highest level of education (> high school) (n,%) | 27 (34.2) | 31 (38.8) |
| Private health insurance (had private health insurance) (n,%) | 6 (7.6) | 9 (11.3) |
| Age (yrs) (mean [SD]) | 56.0 (13.3) | 57.4 (13.6) |
| Height (m) (mean [SD]) | 1.7 (0.1) | 1.7 (0.1) |
| Pain duration (how long have you been troubled with your pain) (yrs) (mean [SD]) | 13.0 (11.9) | 18.5 (15.7) |
| Pain intensity (NRS) (mean [SD]) | 6.7 (1.8) | 6.8 (1.6) |
| Disability score (mean [SD]) | 14.7 (5.2) | 15.8 (5.1) |
| Weight (kg) (mean [SD]) | 91.9 (16.5) | 90.8 (14.6) |
| BMI (mean [SD]) | 32.4 (3.5) | 32.1 (3.6) |
| Utility score (mean [SD]) | 0.6 (0.1) | 0.6 (0.1) |

Abbreviations: yrs years, m metres, NRS Numerical Rating Scale, kg kilograms, BMI Body Mass Index

**Figure 1.**
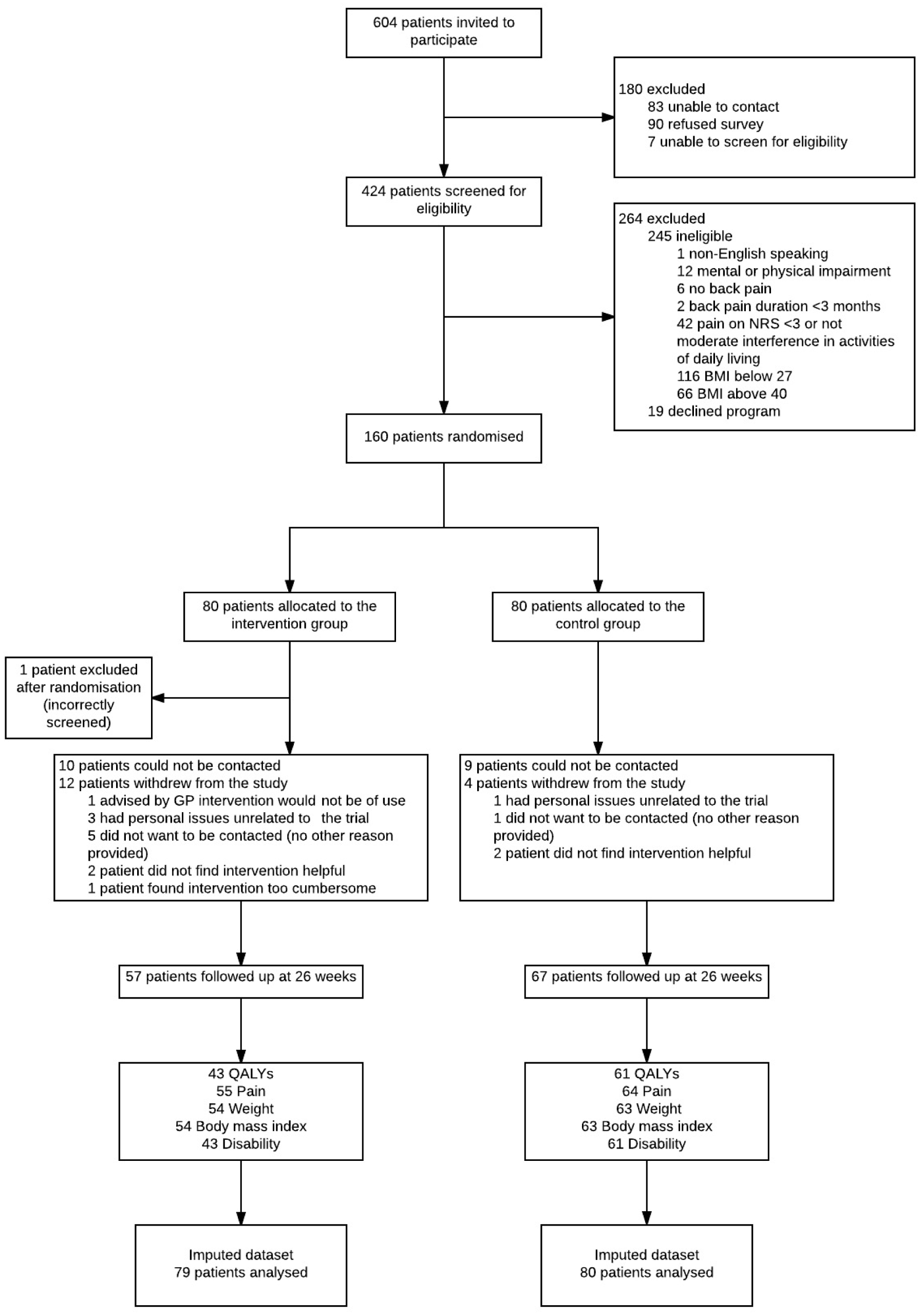
Progress of participants through the study

### Resource use and costs

Of the intervention group patients, 47% (n=37) attended the initial consultation provided by the study physiotherapist and the average number of successful GHS calls was 5.1 (SD 4.5). The mean intervention cost was $708 (SEM 68) per patient. Intervention group participants had significantly lower healthcare costs (-$292; 95%CI: -872 to -33), medication costs (-$30; 95%CI: -65 to -4) and absenteeism costs (-$1000; 95%CI: -3573 to -210) than those of the control group (Table 3). From the societal perspective, the mean total costs were lower in the intervention group than in the control group (-$614; 95%CI: -3133 to 255) (Table 3). From the healthcare perspective, the mean total costs were higher in the intervention group than in the control group ($386; 95%CI: -188 to 688) (Table 2).

**Table 2.** Differences in pooled mean costs and effects (95% Confidence intervals), incremental cost-effectiveness ratios, and the distribution of incremental cost-effect pairs around the quadrants of the cost-effectiveness planes

| Analysis | Sample size | | Effects | $\Delta C$ (95% CI) | $\Delta E$ (95% CI) | ICER | Distribution CE-plane (%) | | | |
| --- | --- | --- | --- | --- | --- | --- | --- | --- | --- | --- |
|  | Int | Cont |  |  |  |  | NE<sup>a</sup> | SE<sup>b</sup> | SW<sup>c</sup> | NW<sup>d</sup> |
| Societal perspective – Primary analysis | 79 | 80 | QALYs | -614 (-3094 to 245) | 0.02 (-0.00 to 0.04) | -31087 | 18.7 | 77.2 | 2.9 | 1.2 |
|  | 79 | 80 | Pain | -614 (-3124 to 243) | -0.35 (-1.33 to 0.64) | 1765 | 15.1 | 61.2 | 18.8 | 4.8 |
|  | 79 | 80 | Disability | -614 (-3133 to 239) | -0.57 (-10.41 to 9.27) | 1087 | 11.0 | 41.5 | 38.5 | 8.9 |
|  | 79 | 80 | Weight | -614 (-3132 to 246) | -2.04 (-4.22 to 0.14) | 302 | 19.3 | 77.6 | 2.4 | 0.7 |
|  | 79 | 80 | BMI | -614 (-3110 to 251) | -0.67 (-1.44 to 0.09) | 915 | 19.3 | 76.5 | 3.5 | 0.7 |
| SA1 – Excluding high absenteeism costs (>\$15,000) | 79 | 79 | QALYs | -8 (-830 to 499) | 0.02 (-0.01 to 0.04) | -432 | 48.4 | 46.2 | 2.5 | 3.0 |
|  | 79 | 79 | Pain | -8 (-837 to 502) | -0.32 (-1.30 to 0.67) | 25 | 38.7 | 35.4 | 13.1 | 12.8 |
|  | 79 | 79 | Disability | -8 (-839 to 504) | -0.32 (-10.23 to 9.59) | 25 | 28.4 | 22.8 | 25.4 | 23.4 |
|  | 79 | 79 | Weight | -8 (-829 to 504) | -2.01 (-4.20 to 0.17) | 4 | 49.8 | 47.0 | 1.6 | 1.5 |
|  | 79 | 79 | BMI | -8 (-830 to 500) | -0.67 (-1.43 to 0.10) | 12 | 49.6 | 46.1 | 2.2 | 2.0 |
| SA2 – Excluding non-adherent participants | 23 | 80 | QALYs | -74 (-2597 to 793) | 0.02 (-0.01 to 0.05) | -3437 | 47.4 | 47.0 | 2.3 | 3.2 |
|  | 23 | 80 | Pain | -74 (-2496 to 800) | -0.83 (-2.09 to 0.42) | 89 | 45.0 | 46.2 | 3.9 | 4.9 |
|  | 23 | 80 | Disability | -74 (-2530 to 787) | -2.92 (-14.24 to 8.39) | 25 | 35.6 | 34.3 | 15.4 | 14.7 |
|  | 23 | 80 | Weight | -74 (-2561 to 793) | -1.33 (-4.19 to 1.52) | 56 | 42.2 | 40.9 | 8.6 | 8.2 |
|  | 23 | 80 | BMI | -74 (-2533 to 794) | -0.43 (-1.38 to 0.52) | 173 | 41.1 | 40.9 | 9.1 | 9.0 |
| Healthcare perspective – Secondary analysis | 79 | 80 | QALYs | 386 (-188 to 688) | 0.02 (-0.00 to 0.05) | 19036 | 91.8 | 4.3 | 0.2 | 3.7 |
|  | 79 | 80 | Pain | 386 (-180 to 691) | -0.37 (-1.35 to 0.60) | -1031 | 74.3 | 3.7 | 0.8 | 21.2 |
|  | 79 | 80 | Disability | 386 (-185 to 687) | -0.88 (-10.78 to 9.03) | -440 | 53.1 | 2.7 | 1.7 | 42.5 |
|  | 79 | 80 | Weight | 386 (-183 to 690) | -2.10 (-4.23 to 0.10) | -187 | 92.8 | 4.3 | 0.1 | 2.8 |
|  | 79 | 80 | BMI | 386 (-176 to 687) | -0.68 (-1.44 to 0.08) | -566 | 91.8 | 4.3 | 0.1 | 3.7 |

Abbreviations: *Int* Intervention, *Cont* Control, *CI* confidence interval, *C* costs, *E* effects, *ICER* incremental cost-effectiveness ratio, *SA* sensitivity analysis.
Note: costs are expressed in 2016 Australian Dollars
<sup>^</sup>All ICERs are for a one point increase in the respective effect measure
<sup>a</sup> The northeast quadrant of the CE plane, indicating that the intervention is more effective and more costly than control
<sup>b</sup> The southeast quadrant of the CE plane, indicating that the intervention is more effective and less costly than control
<sup>c</sup> The northwest quadrant of the CE plane, indicating that the intervention is less effective and more costly than control
<sup>d</sup> The southwest quadrant of the CE plane, indicating that the intervention is less effective and less costly than control

**Table 3.** Mean costs per participant in the intervention and control groups, and unadjusted mean cost differences between study groups during the 26-week follow-up period (based on the imputed dataset)

| Cost category | Intervention group<br>*n*=79; mean (SEM) | Control group<br>*n*=80; mean (SEM) | Mean cost difference<br>(95 % CI) |
| --- | --- | --- | --- |
| Intervention costs | 708 (68) | 0 (NA) | 708 (581 to 850) |
| Healthcare utilisation |  |  |  |
| Healthcare costs | 355 (94) | 648 (175) | -292 (-872 to -33) |
| Medication costs | 119 (12) | 149 (14) | -30 (-65 to -4) |
| Absenteeism costs | 89 (68) | 1089 (652) | -1000 (-3573 to -210) |
| **Total** | **1272 (135)** | **1886 (683)** | **-614 (-3133 to 255)** |

Abbreviations: *n* number, *SEM* standard error of the mean, *CI* confidence interval.
Note: costs are expressed in 2016 Australian Dollars

### Societal perspective: cost-utility

The incremental cost-effectiveness ratios (ICER) for QALYs was -31,087 indicating that one QALY gained was associated with a societal cost saving of $31,087 (Table 2), with 77.2% of the cost-effect pairs located in the south-east quadrant, demonstrating that the intervention was on average less costly and more effective than usual care. The cost-effectiveness acceptability curve (CEAC) for QALYs in Figure 2 (2a) indicates that the probability of the intervention being cost-effective compared with usual care was 0.81 at a willingness-to-pay of $0/QALY, increasing to 0.90 at a willingness-to-pay of $17,000, and reached a maximum of 0.96 at $67,000.

**Figure 2.**
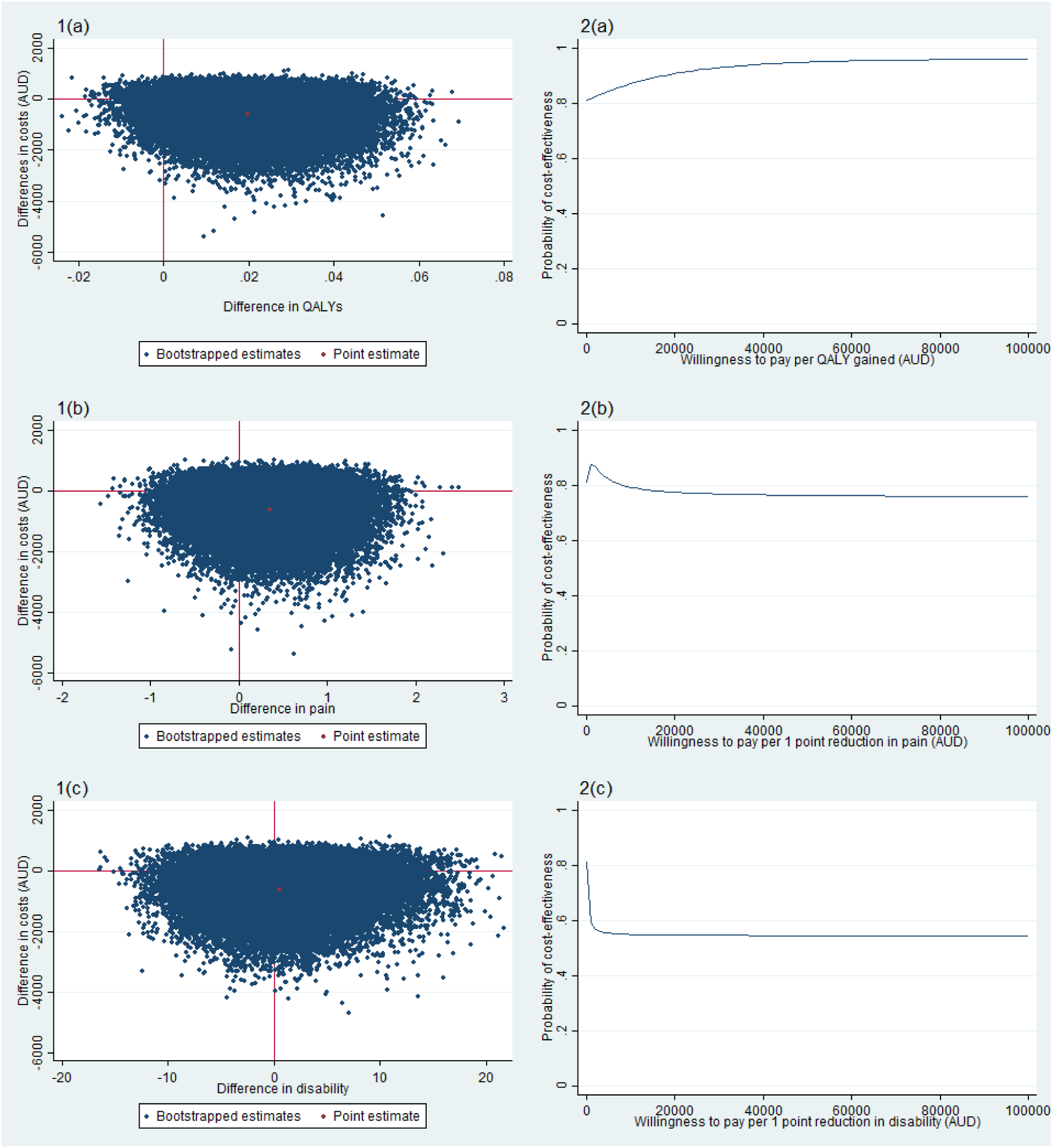
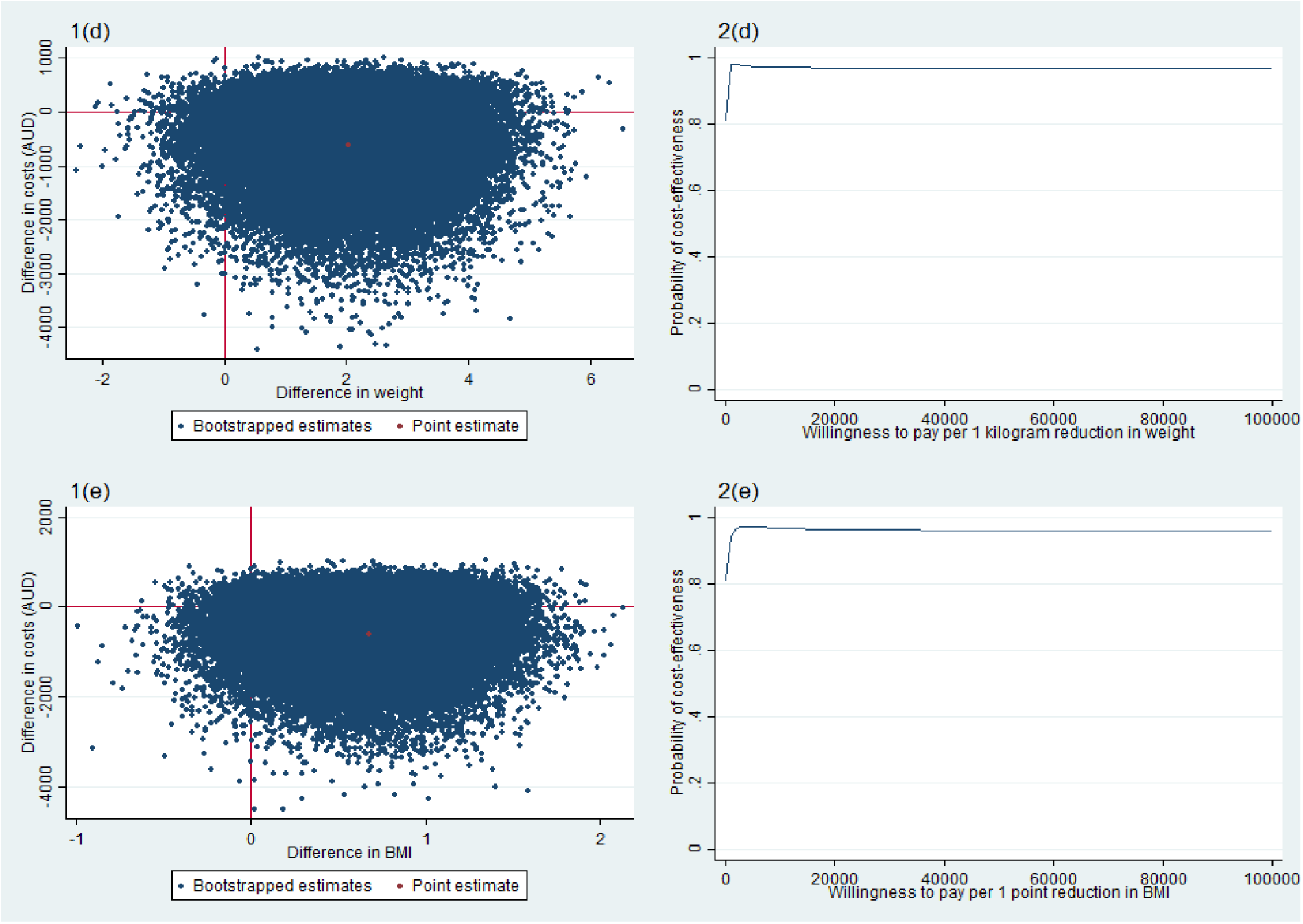
Cost-effectiveness planes indicating the uncertainty around the incremental cost-effectiveness ratios (1) and cost-effectiveness acceptability curves indicating the probability of the intervention being cost-effective at different values ($AUD) of willingness-to-pay per unit of effect gained (2) for QALYs (a), pain (b), disability (c), weight (d) and BMI (e) (based on the imputed dataset).

### Societal perspective: cost-effectiveness

The ICER for pain intensity was 1,765, indicating that a one point decrease in pain intensity was associated with a societal cost saving of $1,765. ICERs in the same direction were found for disability ($1,087 per one point decrease on the Roland Morris scale), weight ($302 per one kilogram weight loss) and BMI ($915 per one BMI point decrease) (Table 2). In all cases, the majority of incremental cost-effect pairs were located in the southeast quadrant (Table 2, Figure 2 [1b-1e]), indicating that the intervention was on average less expensive and more effective than usual care. CEACs for pain intensity, disability, weight, and BMI are presented in Figure 2 (2b-2e).

For all of these outcomes, the probability of cost-effectiveness was 0.81 at a willingness-to-pay of $0/unit of effect. For pain intensity, the probability of cost-effectiveness reached a maximum of 0.88 at a willingness-to-pay of $1000/unit of effect and after this it gradually decreased to 0.76. For disability, the probability of cost-effectiveness decreased with increasing values of willingness-to-pay. For weight and BMI, the probability of cost-effectiveness reached 0.90 at a willingness-to-pay of $1,000/unit of effect (i.e. -1kg or -1 unit of BMI), and remained above 0.90 irrespective of increasing values of willingness-to-pay.

### Societal perspective: sensitivity analyses

The total cost difference between groups was -$8 when we removed one outlier (absenteeism costs > $15,000) from the analysis (SA1), and -$74 when we included only adherent participants (SA2); compared to -$614 in the primary analysis (Table 2).

For QALYs the probability of cost-effectiveness was 0.51 (SA1) and 0.54 (SA2) at a willingness-to-pay of $0/unit of effect. For SA1, the probability of cost-effectiveness increased to 0.90 at a willingness-to-pay of $47,000/QALY, and reached a maximum of 0.92 at a willingness-to-pay of $77,000/QALY. For SA2, the probability of cost-effectiveness increased to 0.90 at a willingness-to-pay of $72,000/QALY, and reached a maximum of 0.91 at a willingness-to-pay of $86,000/QALY. These values are higher than that of the primary analysis (i.e. a probability of 0.90 at a willingness-to-pay of $17,000/QALY).

For pain intensity, the probability of cost-effectiveness was relatively low (i.e. <0.55) at a willingness-to-pay of $0/unit of effect, however, it did reach 0.90 at a willingness-to-pay of $3000/unit of effect in SA2. For disability, in contrast to the primary analysis, the probability of cost-effectiveness remained relatively low (i.e. 0.50 to 0.70) in both sensitivity analyses, regardless of willingness-to-pay. Conversely, for weight and BMI, similar to the primary analysis, the probability of cost-effectiveness reached 0.80-0.90 in both sensitivity analyses.

### Healthcare perspective: cost-utility

For QALYs the ICER was 19,036 indicating that one QALY gained was associated with a cost to the healthcare system of $19,036 (Table 2) and the probability of cost-effectiveness reached a maximum of 0.90 at a willingness-to-pay of $98,000/QALY.

### Healthcare perspective: cost-effectiveness

For pain intensity, the ICER was -1,031, indicating that a one point decrease in pain was associated with a cost of $1,031. ICERs in the same direction were found for disability ($440 per one point decrease on the Roland Morris scale), weight ($187 per one kilogram weight loss) and BMI ($566 per one BMI point decrease) (Table 2). The probability of cost-effectiveness for pain intensity and disability did not reach 0.90 at any value of willingness-to-pay. For pain intensity and disability, the probability of cost effectiveness reached a maximum of 0.77 at $27,000/unit of effect and 0.57 at $8000/unit of effect, respectively. For weight and BMI, the probability of cost-effectiveness was similar to the primary analysis reaching 0.90 at $1000/unit of effect and $3000/unit of effect, respectively.

## Discussion

### Key findings

We found that a healthy lifestyle intervention involving brief telephone advice, offer of a clinical consultation involving detailed education, and referral to a 6-month telephone-based healthy lifestyle coaching service was on average less expensive and more effective than usual care from the societal perspective. For QALYs, the intervention had a high probability (0.81) of cost-effectiveness from the societal perspective at a willingness-to-pay of $0/unit of effect, and increased at higher willingness-to-pay thresholds. However, the probability of cost-effectiveness was not as favourable among sensitivity analyses nor from the healthcare perspective.

### Interpretation of findings

Results of the cost-utility analysis from the societal perspective suggest that the intervention can be considered cost-effective compared with usual care for QALYs. From a probability of cost-effectiveness of 0.81 at a willingness-to-pay of $0/QALY, the probability increased to 0.90 at a willingness-to-pay of $17,000/QALY and reached a maximum of 0.96 at $67,000. The intervention had a high probability (>0.93) of cost-effectiveness at the published Australian ($64,000/QALY) and UK willingness-to-pay thresholds ($34,000-51,000/QALY).^31^

Results of the cost-effectiveness analysis from the societal perspective for pain intensity, disability, weight, and BMI appear favourable. However, because society’s willingness-to-pay per unit of effect gained has not been reported/determined for these outcomes, decisions regarding cost-effectiveness would depend on the willingness-to-pay of decision-makers and the probability of cost-effectiveness that they perceive acceptable. Nonetheless, for all of these outcomes there were relatively high probabilities of cost-effectiveness (i.e. 0.81) at a willingness-to-pay of $0/unit of effect and for all outcomes excluding disability, the probability of cost-effectiveness increased to 0.88 or 0.90 at a willingness-to-pay of $1000/unit of effect.

The two sensitivity analyses indicate that the findings from the societal perspective should be interpreted with caution for QALYs, pain intensity and disability. For QALYs, in contrast to the primary analysis the results of SA2 (i.e. excluding patients without reasonable adherence), the intervention may not be considered cost-effective. The probability of cost-effectiveness was relatively low (<0.55) at a willingness-to-pay of $0/QALY and only reached 0.90 at $72,000/QALY, which is above both the Australian and UK willingness-to-pay thresholds.^31^ For pain intensity in SA2 and for disability in both sensitivity analyses, in contrast to the primary analysis the probability of cost-effectiveness was relatively low (i.e. 0.50 to 0.70), regardless of willingness-to-pay.

We also undertook a secondary analysis from the healthcare perspective, this involved considering intervention, healthcare utilisation and medication costs, but not absenteeism costs. From the healthcare perspective, the intervention may be considered cost-effective for QALYs, weight, and BMI depending on the probability of cost-effectiveness that decision-makers perceive as acceptable. However, the intervention seems not to be cost-effective for pain intensity or disability due to relatively low maximum probabilities of cost-effectiveness (i.e. <0.77).

### Comparison with the literature

This study is the first economic evaluation of a healthy lifestyle intervention for patients with chronic low back pain. As such, direct comparisons to similar interventions are limited. Nonetheless, similar to our findings, other conservative approaches appear to be cost-effective relative to usual general practitioner (GP) care.^32,33^ Specifically, exercise alone or exercise plus GP care and/or spinal manipulation is cost-effective compared to GP care alone; and cognitive behavioural therapy plus physiotherapy is cost-effective compared to GP care alone.^32,34^ However, systematic reviews in this area indicate these results warrant some caution based on overall methodological quality.^32–34^ Our study utilises recommended contemporary methods of economic evaluation and provides comprehensive data to guide decisions about healthcare for this patient group.

### Strengths

A strength of this study is the pragmatic RCT design, meaning the study was completed under ‘real world’ conditions. The design is advantageous for decision-makers to use the study’s findings to guide decisions about real world healthcare services. Another strength of this study is the use of contemporary methods for cost-effectiveness analyses including SUR and bootstrapping. SUR was used to account for potential correlation between cost and effect data and bootstrapping allowed for estimation of uncertainty around the right skewed cost-effectiveness estimates.

### Limitations and directions for future research

A limitation of this study is the amount of incomplete data. The amount of missing outcome data varied between the effect measures however, was at least 25% in all cases. Cost data was missing for 41% of participants after 26-weeks. These levels of missing data are common in economic evaluations of interventions delivered in real-world settings.^35^ We used multiple imputation to account for the missing data, which is recommended over complete case analyses, despite this, results from this study should be treated with caution. A further limitation is that costs were based on participant recall. This may have introduced recall bias, although the period over which participants were required to report their resource use was reasonably short (6 weeks). This study was completed over a relatively short follow-up period of 6 months. It is unknown whether the cost-effectiveness estimates from this study would be similar over a longer follow-up period. Assessing the cost-effectiveness of lifestyle interventions for chronic low back patients over the longer term could possibly produce more meaningful insight. Lastly, the study did not include measures of presenteeism, i.e. reduced productivity while at work. As presenteeism is a potentially significant cost of chronic low back pain,^4^ further research in this area should include such a measure.^36^

### Implications for policy

We found that the intervention group had significantly lower absenteeism and healthcare utilisation costs. These findings suggest that targeting lifestyle risk factors, as part of chronic low back pain management, could result in cost savings from less time off work and reduced healthcare use. Currently, clinical practice guidelines focus on reducing pain and disability, and lifestyle is largely overlooked. Given the global economic burden of chronic low back pain, further recognition of lifestyle as a priority in the treatment of chronic low back pain is warranted. Despite this, inconsistencies among the sensitivity analyses results mean that this interpretation should be treated with caution.

Among other things, decisions to utilise this healthy lifestyle intervention on the basis of cost-effectiveness, would depend on the priority of the decision-maker. This is because we only know how much society is willing to pay per QALY gained, but not for pain intensity, disability, weight, or BMI. Once a decision-maker determines what they value as an outcome, the methodological limitations and variability found in the sensitivity analyses should be considered in the decision to utilise this intervention. Nonetheless, considering the high prevalence of chronic low back pain globally, and limited resources available to support such patients, this study provides decision-makers with valuable information to guide decisions about the utility of available interventions.

## Conclusions

We conducted an economic evaluation of a healthy lifestyle intervention involving brief telephone advice, offer of a clinical consultation involving detailed education, and referral to a 6-month telephone-based healthy lifestyle coaching service for patients with chronic low back pain, who are overweight or obese. The intervention seems to be cost-effective for QALYs from the societal perspective but not from the healthcare perspective. Variability found in the sensitivity analyses findings should be considered in the decision to utilise this intervention.

